# Altruistic disease signalling in ant colonies

**DOI:** 10.1101/2024.02.27.582277

**Authors:** Erika H. Dawson, Niklas Kampleitner, Jennifer Robb, Florian Strahodinsky, Anna V. Grasse, Sylvia Cremer

**Affiliations:** ISTA (Institute of Science and Technology Austria), 3400 Klosterneuburg, Austria; Université Sorbonne Paris Nord, 93430 Villetaneuse, France; Technical University Munich, 81675 Munich, Germany

**Keywords:** host-parasite interactions, chemical communication, social immunity, disease signals, honest signalling, cooperation, ants

## Abstract

Sick individuals often conceal their disease status to group members, thereby preventing social exclusion or aggression. Here, we show that infected ant pupae, on the contrary, actively emit a chemical signal that triggers their own destruction by colony members. In our experiments, this altruistic disease-signalling was performed only by worker but not queen pupae, reflecting differences in their immune capabilities, as worker pupae suffered from extensive pathogen replication whereas queen pupae were able to restrain infection. Inducing others to sacrifice oneself, only if one’s own immunity fails, suggests a fine-tuned interplay between individual and social immunity, efficiently achieving whole-colony health.

## Main text

Sick animals living in groups are frequently excluded or aggressed by other group members in order to prevent cross-contamination, but also to take advantage of the weak status of the infected individual^1,2^. As a result, these individuals often conceal their infection status by, for example, suppressing sickness behaviours in the presence of conspecifics. This is not the case when group members are kin, since relatedness diminishes the conflict of interest between sick and healthy individuals. Therefore, even when group members can detect disease in others, for example using sickness cues such as altered physical appearance or behaviour, avoidance is often only displayed against non-kin, whilst normal social interactions are maintained with diseased kin^3^.

Diseased individuals can therefore benefit, not only by not concealing, but even from actively communicating their health state to their relatives in order to receive additional care. As such, fungal pathogen-exposed termites display a vibratory signal inducing grooming behaviour by nestmates and wounded ants produce behavioural and chemical displays attracting their colony members to care for their wounds, which in both cases reduces the signalling individual’s infection risk and hence improves its survival^4,5^. Helping a relative to survive infection can also indirectly benefit the caregiver, by increasing own inclusive fitness via shared genes with the care-receiving individual^6^. Active disease-state signalling is therefore selected for in groups of relatives, when both the signaller and the signal-receiver gain from the care response. But what happens when the related group member’s reaction is not care but instead exclusion or aggression towards the infected individual, to protect the group? Should the infected individual still signal its sickness to others, thereby risking its own sacrifice for the group’s benefit?

The evolution of such ‘altruistic disease signalling’ in social groups should be promoted by two factors: (i) high relatedness, leading to a large indirect fitness gain to the sacrificed individual, if signalling promotes group health, and (ii) low direct fitness loss to the sacrificed signaller, i.e., when its expected future reproductive value is small. Both are fulfilled in the non-reproductive workers of social insect colonies. Workers are typically highly related with one another and produce no offspring of their own, but instead rear the queen’s brood and maintain the colony. Therefore, the fitness of workers depends on the fate of the colony as a whole. Here, we experimentally test whether altruistic signalling of one’s own sickness indeed evolved in social insect colonies.

Social insects, particularly termites, ants and social bees, show sophisticated collective disease defences when colony members come into contact with pathogens. These social immunity measures range from sanitary care of individuals that can still be rescued, to removal of fatally-infected colony members^7,8^. Here we studied the invasive garden ant, *Lasius neglectus*, in which adult workers prematurely unpack worker pupae with deadly infections of the fungal pathogen *Metarhizium brunneum*, then destroy and disinfect them, thereby preventing pathogen replication in the host and ultimately spread throughout the colony^9^. The expression of this behaviour relies on the detection of a disease-related change of the pupal surface chemistry. In particular, unpacked pupae showed increased abundance of four cuticular hydrocarbon (CHC) peaks (C33:2, C33:1, C35:2, C35:2+C35:1), and experimental CHC removal prevented unpacking behaviour^9^. Yet, it remains unknown whether these chemical changes are simply passively-emitted cues resulting from infection or the induced immune response, or whether they represent active signalling by the infected pupae to the workers to trigger their own destruction for the benefit of the colony, thereby increasing their own inclusive fitness. In order to disentangle these two scenarios, we manipulated the context in which pupae could signal their disease status.

Individual pupae were experimentally infected with the pathogen, or sham-treated, and then kept either with two workers, or alone. We employed this full-factorial approach as we speculated that if a signal is costly to produce (e.g.^10^) and pupae are able to sense the presence or absence of workers, then they should only signal when both infected and in the presence of the intended receivers, i.e. the workers. Since the close social interactions in ant colonies often involve the exchange of substances, including the ants’ own CHCs^11^ we needed to disentangle pupae- and worker-derived chemical compounds to be able to exclusively quantify the pupa-produced CHCs. To this end, we used stable isotope labelling to enrich only the workers’ but not the pupal CHCs with the natural carbon isotope ^13^C by feeding the workers ^13^C glucose ahead of the experiment (as detailed in the Online Methods; Fig. S1). We focused our analysis on the four previously identified candidate peaks for possible unpacking cues/signals, with two of them (tritriacontadiene, C33:2, and tritriacontene, C33:1) in particular being associated with immune system activation^9^.

Our chemical analysis clearly revealed that only the two immune-associated compounds were upregulated, and only by pupae that were both infected and kept with workers (Fig. S2A; [LM] infection treatment*worker presence interaction: C33:2: LR χ^2^=7.044, df=1, p=0.008, C33:1: LR χ^2^=4.419, df=1, p=0.0356; Table S1a), so that their chemical profile deviated from that of the other three groups (Fig. 1A, Table S2; all posthoc p-values to the other three groups <0.05). The fact that only the two immune-associated compounds were increased in abundance by infected pupae with workers, and that infection load did not predict abundance of either compound (both p>0.16; Table S1b), provides double evidence that this upregulation is not directly brought about by the fungus itself, but is instead under dynamic host control, allowing the pupae to actively signal their disease status. Interestingly, these two same unsaturated hydrocarbons also show increased levels in virus-infected (C33:2)^12^ and bacterial cell wall immune-stimulated (C33:1)^13^ honeybees, and C33:1 was found to induce bee hygienic behaviour^14^. The same compounds described here to be actively upregulated for disease-signalling during fungal infection in ants are therefore at least also passive cues of viral and bacterial disease in a second representative of the social Hymenoptera (ants, bees and wasps). Conversely, different compounds are modulated upon *Metarhizium* fungal infection in the unrelated second group of social insects, the termites^15^. More studies on active signalling across social host-pathogen systems are needed to evaluate the intriguing hypothesis emerging from these findings of the possible existence of a conserved universal disease signal across the social Hymenoptera with an important role in the evolution of altruistic disease signalling.

**Figure 1.**
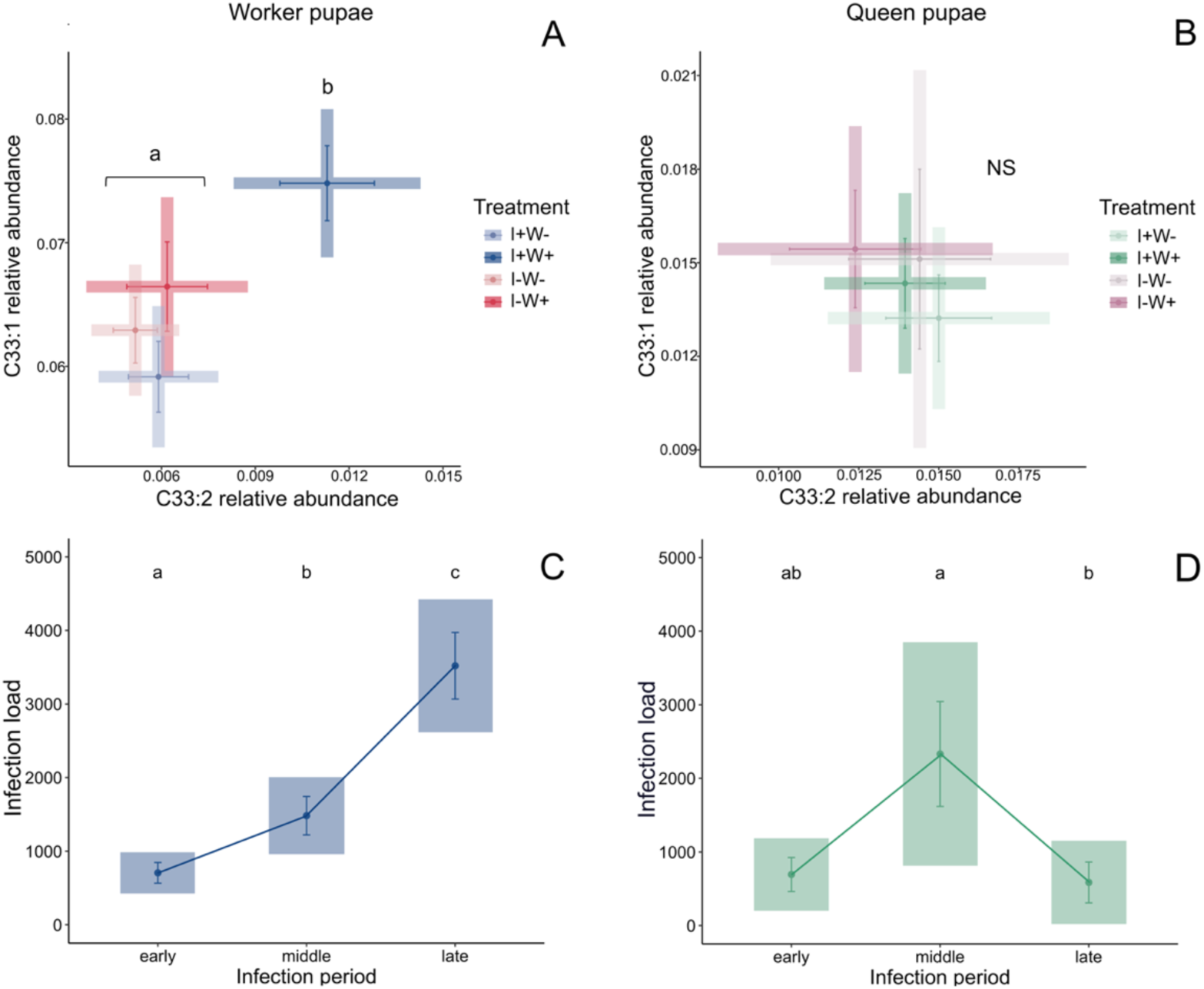
Chemical signalling and infection load of infected ant pupae. (A) *L. neglectus* worker pupae carrying an infection with *M. brunneum* (I+, blue) showed an upregulation of the relative abundance of the two immune-associated cuticular hydrocarbons tritriacontadiene, C33:2, and tritriacontene, C33:1, in the presence of tending workers (W+; dark tone) only, so that they differed in their chemical profile from the infected pupae in the absence of workers (W-; pale tone), as well as the pathogen-free control pupae (I-; pink) independent of worker presence (Tables S1,S2). (B) Queen pupae, in contrast, showed no difference between the four treatment groups (infection green, sham purple; statistics: Tables S3,S4). The pupal infection load (given as fold-change of its measured fungal load to the maximum caste-specific exposure dose) of (C) worker pupae increased steadily over time, whilst in queen pupae (D) infection was reduced back to early infection levels over the course of the experiment (Table S7). Graphs show means ± sem as dots and whiskers in shaded 95% CI box. Letters denote significant posthoc two-sided p<0.05 after correction for multiple testing; ns: non-significant. Chemical analysis (A,B) based on a total of 426 pupae, infection load analysis (C,D) on 265 infected pupae.

While altruistic disease signalling is expected to evolve in sterile workers, the situation is more complex for the daughter queens. By alerting others to destroy them, queen pupae would risk losing potential future reproduction if they would survive infection. On the other hand, by spreading their infection to their colony, they could incur high indirect fitness costs. We therefore tested if infected *L. neglectus* queen pupae showed similar altruistic disease signalling as worker pupae, or not. We found that – contrary to the worker pupae – none of the four candidate compounds (Figs. 1B, S2B; [LM] all p-values >0.8; Table S3a) nor any of their other compounds (Online Methods, Table S5 [LM] all p-values >0.14) showed a difference between the four treatment groups. We could therefore not detect any disease signalling in the queen pupae, despite the queen pupae also clearly developing an infection (Online Methods; Fig. 1D). Consequently, whilst the signalling infected worker pupae elicited unpacking above baseline levels (Cox proportional-hazards regression, treatment effect p=0.0037; Table S6), this was not the case for the non-signalling infected queen pupae (p=0.115; Table S6), meaning they were not targeted for destructive disinfection by the workers.

We asked if the absence of signalling in the queen pupae represents a selfish behaviour which puts the colony at risk, or an honest representation of the higher immunocompetence of the queen caste in social insects^16,17^, allowing them to cope with infection and making the need for signalling obsolete. To this end, we compared pathogen load progression in queen and worker pupae. We found that, whilst infection load steadily increased over the course of infection – with no sign of even plateauing – in the worker pupae (Fig. 1C; all p-values between early, middle and late infection period p<0.0158; Table S7a), queen pupal infection load peaked intermediately, but then decreased by almost 3-fold (Fig. 1D; middle to late infection period p=0.006, Table S7b). This suggests that, since the more potent queen immune system was able to reduce infection, whilst the feebler worker immune system was not, both the signalling in the worker pupae and the absence thereof in the queen pupae are honest reflections of the caste-specific differences in individual infection progression during our experiment. Future studies could identify under what infection conditions individual immunity of the queen pupae may fail and whether this would lead to signalling also in the queen caste.

The observed expression of altruistic disease signalling only in pupae with an overwhelmed immune system suggests a fine-tuned balance between individual and social immunity in social insect colonies. Universal destruction of all infected colony members based on passive cues of sickness would instead disregard varying levels of immunity and recovery capabilities, potentially causing substantial, avoidable losses, including costly-to-rear individuals like new queens. Such mechanisms of individual health assessment, and calling others to destroy oneself only when individual immunity fails, instead allows concurrent protection of the individual colony members and overall colony health.

## Methods

### Ant host

As host species, we used the invasive garden ant, *Lasius neglectus*. We collected several hundred workers, multiple queens and brood of this species from its introduced supercolonial populations^18,19^ in Jena, Germany (N 50° 55’ 54.599” E 11°35’ 8.401”) in July 2018, August 2019 and June 2021 and in Seva, Spain (N 41°48’ 32.699" E 2° 15’ 42.3") in April 2016 and May 2018. Collection of this unprotected species from the field was performed in compliance with international regulations, such as the Convention on Biological Diversity and the Nagoya Protocol on Access and Benefit-Sharing (ABS; permit numbers: ESNC12 and SF/0558-0561). After collection, ants were reared in large stock colonies in the laboratory with 30% sugar water and minced cockroaches. Experiments were performed with pupae of a standardized developmental stage (white pupae with black eyes) and with workers, always sampled from the same nest as the respective pupae from the inside of the brood chamber to avoid foragers. All experimental work followed European and Austrian law and institutional ethical guidelines.

### Pathogen infection

As a pathogen, we used the entomopathogenic fungus *Metarhizium,* whose infectious conidiospores naturally infect ants^20–22^ by penetrating their cuticle, killing them and growing out to produce highly-infectious sporulating cadavers^23,24^. We infected the pupae with conidiospores (abbreviated as ‘spores’ throughout) of *M. brunneum* strain Ma275 (KVL 04-57, obtained from J. Eilenberg and N.V. Meyling from the University of Copenhagen, Denmark), which were freshly cultivated from long-term storage on Sabouraud 4% Dextrose Agar plates (SDA; Sigma-Aldrich) at 23°C and had been harvested in sterile Triton X-100 (Sigma-Aldrich; 0.05% in Milli-Q water). Spore germination was confirmed to be above >98% one day prior to each experiment.

We exposed each pupa individually to the fungal pathogen by placing it on a glass slide, applying the fungal suspension and rolling the pupa in the suspension using soft forceps. Control pupae were treated the same, except for using Triton X-100 solution as a sham treatment. Pupae were then left to air-dry on the slide before putting them in a plastered dish (90×35 mm, Licefa GmbH) for three days before the start of the experiment. We used 1 µl of a 1 x 10^6^ spores/ml fungal suspension for the exposure of worker pupae and 2 µl of a 1 x 10^7^ spores/ml fungal suspension for queen pupae, and the same respective amounts of sterile 0.05% Triton X-100 (Sigma-Aldrich; 0.05% in Milli-Q water) for the control pupae. Using the below-detailed fungal load determination by quantitative real-time PCR (qPCR), we determined that this exposure procedure resulted in an approximately 10-fold higher exposure dose for the queen pupae than the worker pupae (queen pupae: mean 1.587 x 10^-3^ ng/µl, maximum 2.232 x 10^-3^ ng/µl, n=3; worker pupae: mean 1.603 x 10^-4^ ng/µl, maximum 2.158 x 10^-4^ ng/µl, n=7). We exposed the queen pupae to this higher spore load, since they are much larger in size than the worker pupae (mean queen pupa length: 5.7 mm, width: 2.5 mm; n=4; mean worker pupa length: 2.9 mm, width: 1.1 mm; n=4, leading to an approx. 4-fold larger surface and 9-fold higher volume of queen pupae compared to worker pupae) and preliminary data revealed that an approx. half concentration of the final dose was insufficient to induce a successful infection in the queen pupae. This is in line with reports that, generally, social insect queens have higher immunocompetence than workers^17^ and, in particular, that *Lasius* queens show higher immune gene expression than workers^16^.

To allow for successful establishment of infection, pupae were kept in plastered dishes (90×35 mm, Licefa GmbH), after exposure, for three days in the absence of workers (as in^9^), as otherwise the sanitary care provided by accompanying workers would interfere with the infection process due to spore removal and disinfection^25,26^. Equally, control pupae were kept without workers for the same duration after their exposure to the sham treatment to ensure they were the same age as the fungus-treated pupae. This establishment phase and all following experimental procedures took place in a temperature and humidity-controlled room at 23°C, 65% RH and a 12h day/night light cycle and the plastered dishes were watered every 2-3 days to keep humidity high.

### Experimental procedures

After the three days of fungal (or sham) infection-establishment, individual pupae were placed into a plastered dish (35×10mm; Falcon or SPL Life Sciences), either alone or with two workers that originated from the same colony as the pupa. Using a full-factorial design, we thereby set up four treatment groups: infected pupae without workers (I+W-), infected pupae with workers (I+W+), control pupae without workers (I-W-) and control pupae with workers (I-W+). From each treatment, we sampled pupae over a period of 42h after setting them up with the workers, at seven time points in intervals of six hours (i.e. at 6, 12, 18, 24, 30, 36 and 42 hours), to prevent them from being damaged by the destructive disinfection by the workers. In the W+ groups, we noted if the pupa was unpacked or not, but excluded pupae that appeared themselves damaged or dead, to ensure we only analysed pupae that were still able to modulate their cuticular hydrocarbons (CHCs). At each sampling time point, we also sampled control pupae. Furthermore, we only included replicates, in which both workers were still alive at the end of the sampling period. Lastly, a few samples had to be excluded after chemical analysis as they did not match our quality criteria as detailed below. Our final sample size therefore consisted of 426 pupal samples (323 worker pupae, of which 69 I+W-, 133 I+W+, 64 I-W-, 57 I-W+, from both the Jena (n=168) and the Seva (n=155) populations; 103 queen pupae, of which 19 I+W-, 45 I+W+, 19 I-W-, 20 I-W+, all from the Jena population). We further used a total of 510 workers in our experiments (380 with the worker pupae and 130 with the queen pupae). We froze each pupa with its cocoon (as in Pull et al. 2018^9^) in a brown glass vial (1.5 mL, Agilent) closed with one-component-closure-caps (Agilent) at -80°C, to store them for later chemical extraction for determination of the pupal surface chemicals, followed by DNA extraction and quantification of the fungal load. For the replicates containing workers, we pooled both workers in one glass vial for later chemical analysis (detailed below). We conducted the experiment separately for queen and worker pupae, with each experiment lasting five days, from the time of exposure, over the three days of isolation to the last sampling point, each carried out in a single block.

### Chemical analysis

#### Gas chromatography–mass spectrometry

We used gas chromatography–mass spectrometry (GC–MS) to determine the composition of CHCs of worker and queen pupae, as well as their attending workers (where appropriate, i.e. in I+W+ and I-W+). To this end, we extracted pupal compounds from individual worker and queen pupae, as well as the worker compounds from the pool of both workers per replicate, by adding 90 µL *n*-pentane solvent (Supelco) to the vials in which they had been collected. Vials were closed with aluminium-faced silicon septa caps (Agilent). Extractions were performed with the solvent for 5 min under gentle agitation at room temperature. The supernatant was transferred to glass vials with 350 µL glass inserts and sealed with aluminium-faced silicon septa (Agilent). The *n*-pentane solvent contained two internal standards at the beginning and the end of the *L. neglectus’* range of hydrocarbons (C27 - C37), *n*-tetracosane and *n*-hexatriacontane at 0.1 µg/mL concentration (both CDN Isotopes), both fully deuterated to enable spectral traceability and separation of internal standards from ant-derived substances. The extracts from the different treatment groups were run in a randomized manner, intermingled with blank runs (containing only *n*-pentane), and handling controls, using GC-MS (gas chromatograph GC7890 coupled to mass spectrometer MS5975C [Agilent Technologies, Santa Clara, California]).

A liner with one restriction ring filled with borosilicate wool (Joint Analytical Systems) was installed in the programmed temperature vaporization (PTV) injection port of the GC, which was pre-cooled to -20°C and set to solvent vent mode. 50 µL of the sample extractions were injected automatically into the PTV port at 40 ml/s using an autosampler (CTC Analytics, PAL COMBI-xt, CHRONOS 4.2 software [Axel Semrau]) equipped with a 100 µL syringe. Directly after injection, the temperature of the PTV port was increased to 300°C at 450°C/min, thereby vaporizing the sample analytes and transferring them to the column (DB-5ms UI; 30 m × 0.25 mm, 0.25 μm film thickness [Agilent]) at a flow rate of 1 ml/min. To ensure optimal peak separation and shape, helium was used as the carrier gas at a constant flow rate of 3.0 ml/min for 2 min, then decreased to a constant flow rate of 1.1 ml/min at 100 ml/min^2^ and the oven was programmed to hold 35°C for 4.5 min, then ramp to 325°C at 20°C/min, and hold this temperature for 10 min. The GC-MS transfer line was set to 325°C, and the mass spectrometer (MS) operated in alternating TIC-SIM mode with a scan range of 35-600 amu in the total ion current (TIC) mode electron ionization mode (70 eV; ion source 230°C; quadrupole 150°C, with a detection threshold of 150). Disentanglement of worker-applied and pupa-produced compounds (using the below detailed quantification of ^12^C and ^13^C) was based on the SIM (selected ion monitoring) mode, where the ions 57.1, 58.1, 59.1, 60.1, and 61.1 were selected. In addition, a C7-C40 saturated alkane mixture (in steps of 1 at concentration of 0.1 µg/mL for each alkane in pentane; Supelco) was run as external standard, enabling calculation of Kováts retention indices (RIs) and correcting for possible shifts in retention time during the time the samples were run. Compound peaks were extracted from the total ion current chromatograms (TICCs) of representative sample runs, using a deconvolution algorithm (MassHunter Workstation, Qualitative Analysis B.07.00 [Agilent Technologies]). Compound identification was performed by comparing the mass spectra and Kováts index of each compound to the Wiley 9th edition/NIST 11 combined mass spectral database (National Institute of Standards and Technologies), by manual interpretation of diagnostic ions, as well as by comparison to previous cuticular hydrocarbon (CHC) compounds identified for *L. neglectus* adult workers^18^ and worker pupae^9^. For reasons detailed below, this study does not include all previously described CHCs, only the ones which fitted our criteria to determine ^13^C levels accurately in the method we established here to be able to disentangle worker-applied from pupa-produced CHC production (detailed below). Table S8 lists all CHCs used in this study.

#### 13C-enrichment in worker-produced CHCs

In this study, we wanted to compare the pupal compound production in dependence of both their infection status (infected I+ vs uninfected I-) and the presence or absence of attending workers (W+ vs W-). As it is known that colony members exchange CHCs to form a colony “gestalt” odour^11,27^, we developed a method to quantify exclusively the pupa-produced subset of the CHCs found on the pupae, removing any possible worker-applied compounds, to be able to determine pupal signalling. To do this we used stable isotope labelling^28,29^ of the worker compounds by feeding workers glucose enriched with ^13^C, which is a stable natural isotope to ^12^C, making up 1.1% of environmental carbon. By feeding workers glucose containing a higher than natural ^13^C content (99% than the natural 1.1%), ^13^C becomes integrated into the workers’ CHC compounds at enriched levels compared to the natural levels. When letting these ^13^C-enriched workers tend normal pupae (i.e. pupae from stock colonies reared on normal, non-^13^C-enriched glucose), the workers transfer their ^13^C to the pupae, altering the natural ^12^C to ^13^C ratio in the extracts obtained from the pupae (see below). Quantification of the ^12^C to ^13^C ratio of each compound of (i) the tending workers (to determine the exact ^13^C integration level into the workers’ CHCs for each specific compound and replicate) and (ii) the pupa kept with these workers, will therefore indicate which proportion of the CHC is from non-enriched pupal origin and which from enriched worker origin. Excluding the worker-applied proportion by use of the below-detailed formula thus allowed us to calculate the quantity per compound produced exclusively by each pupa in response to its infection status and the social situation (presence or absence of workers) it was subjected to.

In detail, worker ants were put in experimental containers (90×35 mm, Licefa GmbH) in groups of 50 (worker pupae experiment: n=20 groups; queen pupae experiment: n=11 groups) and fed exclusively with ^13^C-enriched glucose solution (D-Glucose-^13^C_6_, 99 atom % [Eurisotop], 10% weight/volume in water) for three weeks prior to being placed with the pupae in the experiment. Throughout these three weeks, the pupae, later used in the experiment, remained in the stock colonies and had only ever received normal, non-labelled glucose. After these three weeks, two workers from the same group, were put with an individual pupa originating from the same stock colony, as detailed in the experimental protocol above. Besides sampling the pupae in the six-hour intervals described above, we also sampled their two accompanying workers to analyse their CHCs by the above-mentioned GC-MS procedures, using automatized integration of the peak area of each compound and standardization of the amount of each peak to its closest internal standard eluting before (MassHunter Workstation, Quantitative Analysis B.07.00 [Agilent Technologies]), thereby correcting for possible injection differences. For each compound, we quantified the ^12^C and ^13^C proportion based on the SIM scans, using the generally highly abundant ion 57.1 as quantifier ion for ^12^C (as it represents the fragment ^12^C_4_H_9_), and the sum of ions 58.1, 59.1, 60.1 and 61.1 as quantifier ions for ^13^C, as they reflect the level of ^13^C enrichment (^13^C_1_^12^C_3_H_9_, ^13^C_2_^12^C_2_H_9_, ^13^C_3_^12^C_1_H_9_ and ^13^C_4_H_9_ respectively; simplified scheme due to low natural abundance of other possible isotopes). Since ^13^C makes up 1.1% of environmental carbon, its natural level in C_4_H_9_ – and hence the baseline found in non-enriched samples (Fig. S1) – is 4.4%.

Using this method, we found that the CHC profiles of the workers fed with ^13^C-labelled glucose do indeed show compound-specific ^13^C levels increased above the natural values of 4.4% and that the workers transfer their CHCs to pupae during their social contact with the pupae: pupae kept with workers had increased ^13^C levels > 4.4%, whilst this was not the case in pupae kept without workers, which was equally true for worker (Fig. S1A) and queen (Fig. S1B) pupae. Notably, the intensity of ^13^C-enrichment differed between worker groups (likely depending on differences in feeding activity), yet the difference between individual workers within feeding groups was negligible (as workers of the same feeding group differed only by a mean of 2.2% in their proportion of ^13^C in compound C33:2 and 3.1% in C33:1, as determined by quantification of n=19 additional pairs of workers from 4 feeding groups). To determine the level of ^13^C integration for the two workers per replicate, we therefore pooled both workers for a joint extraction in the experiment to achieve the replicate-specific worker ^12^C and ^13^C for each of their CHC compounds, which was required to then determine the pupae-produced quantity for each compound of each pupa.

#### Subtraction of worker-transferred CHCs from total pupal profiles

^13^C labelling of worker-derived CHCs allowed us to subtract the transferred worker CHCs from the total compound amount measured in the pupae (pupa- and worker-derived). This enabled us to quantify the CHCs produced exclusively by the pupae, which we then compared between treatments. As only the workers but not the pupae had been ^13^C-enriched at the start of the experiment, the proportion of ^13^C in solely pupa-produced compounds reflected the natural 4.4% in the four C atoms (as also confirmed by pupal CHCs in the absence of workers, Fig. S1). Consequently, any measured value above 4.4% of ^13^C compared to ^12^C in any compound, must reflect application of the ^13^C-enriched worker CHCs. Since we measured for each compound and replicate, how much ^13^C had been integrated in the worker CHC due to the feeding of labelled glucose, we knew both, the proportion of ^13^C for the workers (as quantified from the experiment) and from the pupa (natural 4.4%). Given the measured value for total (pupa- and worker-derived) ^12^C and ^13^C per compound, we could therefore disentangle the pupa- and worker-derived contribution and reach the purely pupa-produced quantity (A_pupa_) for each compound by the following formula:

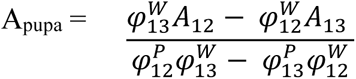

Where, for each compound:

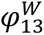 or 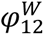 = the proportion of ^13^C or ^12^C of the compound measured in the accompanying workers (quantified from the replicate-specific worker extract)

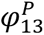 or 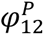 = the natural proportion of ^13^C or ^12^C in the unenriched pupal CHC (set to 0.044 and 0.956 respectively)

A_12_ = the total amount of ^12^C of the compound quantified from the pupal extract (reflecting both pupa- and worker-derived ^12^C)

A_13_ = the total amount of ^13^C of the compound quantified from the pupal extract (reflecting both pupa- and worker-derived ^13^C)

Note that this calculation was performed for each compound and replicate, using the amounts obtained by standardisation to the closest internal standard eluting before the compound (as detailed above). From the previously described CHCs of *L. neglectus* worker pupae^9^, five (3-methylhentriacontane; 13,23-dimethylpentatriacontane; 11,25-dimethyl– pentatriacontane; 7, 11, 23-trimethylpentatriacontane; *n*-hexatriacontane) could not be included in the current analysis, as they were either coeluting with other compounds impacting quantification or ^13^C was undetectable in many of the pupal and worker samples (note that four of these excluded compounds were analyzable in the queen pupae due to higher compound abundance, yet their inclusion did not alter our findings so for consistency reasons we report on the compounds that are conclusive for both worker and queen pupae). Worker and queen pupae shared the same compounds (Table S8).

Furthermore, we could not include 35 worker and one queen pupae samples, which had been originally sampled, but for which accurate assessment of their pupa-produced CHC quantity was not possible for all peaks (these pupae are already deducted from the final sample sizes reported above). This was either because – likely due to natural variation or measurement inaccuracies at low compound abundances – the ^13^C proportion of some of their compounds had a higher value than that of accompanying workers (worker pupae: n=21; queen pupae: n=0), or as their ^13^C was below detection threshold in some compounds, which were typically overall of low quantity (worker pupae: n=14; queen pupae: n=1). When the measured ^13^C proportion of a compound was smaller than 4.4% (similarly likely due to natural variation around the 4.4% average or measurement inaccuracies), which occurred in 10.68% of the worker pupae compounds (621 out of 5814) and 18.39% of the queen pupae compounds (341 out of 1854), we used the measured value as pupa-produced amount because workers cannot have transferred said compound. Similarly, when no detectable amount of a compound in either both ^12^C and ^13^C, or only in ^13^C occurred in the accompanying workers (in 134 cases in the worker pupae and 85 cases in the queen pupae), or when workers were absent from the treatment group (i.e. in all I+W- and I-W-groups), the measured total value of the pupa was set as the pupa-produced amount. For each pupa-produced compound, we then calculated its relative proportion in the overall pupal bouquet and report these relative abundances (standardised amount of the respective compound / sum of the standardised amounts of all 18 compounds). Our main focus was to see whether the relative abundances of the four CHCs that were identified by Pull et al. 2018^9^ as possible candidate compounds for either a cue or signal were differentially modulated by the pupae in response to treatment.

### Pathogen load quantification

We used real time PCR to quantify the ‘fungal load’ of all fungus-treated worker and queen pupae sampled from the experiment, with the exception of one of the worker pupae samples, which was lost during DNA extraction, leading to a final sample size of n=201 worker and n=64 queen pupae. We also analysed all sham-treated pupae (except for three samples that could not be analysed due to technical issues; leading to 119 control worker pupae and 38 control queen pupae), and used their highest value to set the concentration threshold above which we could confidently identify fungal infection and perform an accurate fungal load quantification using real time PCR (qPCR), quantifying absolute fungal DNA load using the standard curve method (detailed below). With a value of 7.2 x 10^-4^ ng/µl this threshold value was approx. 500-fold and 3000-fold lower than the mean fungal load of the worker pupae (3.92 x 10^-1^ ng/µl) and queen pupae (2.34 ng/µl) respectively. We also quantified the fungal load of seven worker and three queen pupae samples frozen straight after exposure, to obtain the worker pupae and queen pupae-specific ‘exposure dose’ (reported above). As these exposure doses differed between the castes, we calculated the ‘infection load’ for each pupa as its measured fungal load relative to the maximum caste-specific exposure dose to determine the infection progression after exposure.

#### DNA extraction and qPCR

We extracted and quantified *Metarhizium* DNA per infected ant pupae by targeting the sequence of the fungal ITS2 gene using quantitative real-time PCR (qPCR) as in Giehr et al. 2017^30^. Prior to DNA extraction, the samples were homogenized using a TissueLyser II (Qiagen) with a mixture of 2.8 mm ceramic (VWR), 1 mm zirconia (BioSpec Products) and approx. 100 mg glass beads (425-600 µm, Sigma-Aldrich). Homogenisation was carried out in two steps (2 x 2 min at 30 Hz). DNA extractions were performed using the DNeasy 96 Blood & Tissue Kit (Qiagen), following the manufacturer’s instructions, with a final elution volume of 50 μl Buffer-AE. qPCR was performed using primers targeting the *Metarhizium brunneum* ITS2 rRNA gene region (Met-ITS2-F: 5’-CCCTGTGGACTTGGTGTTG, Met-ITS2-R: 5’-GCTCCTGTTGCGAGTGTTTT; Gier et al. 2017^30^). Reactions were performed in 20 μl volumes including 1x KAPA SYBR® FAST qPCR Master Mix (Kapa Biosystems), 3 pmol of each primer (Sigma-Aldrich) and 2 μl of extracted DNA. The amplification program was initiated with a first step at 95°C for 5 min, followed by 40 cycles of 10 s at 95°C and 30 s at 64°C. Quantification was done based on the standard curve method, using standards covering a range from 10^-1^ to 10^-5^ ng/µl fungal DNA (measured using a NanoDrop spectrophotometer, Thermofisher). Each run included the standards and a negative control. Samples as well as standards and controls were run in triplicates. Specificity was confirmed by performing a melting curve analysis after each run.

### Statistical data analysis

Statistical analyses were performed using R Studio v1.4.1103. For all models we checked the necessary assumptions by viewing histograms of data, plotting the distribution of model residuals, checking for unequal variances and the presence of multicollinearity and assessing models for influential observations. We transformed the data to obtain normality, using the package ‘bestNormalize’^31^ (using either order-norm, standardized box-cox, standardized log, standardized square-root, Yeo-Johnson, or asin transformations). Figures show the untransformed data.

For all models, we first tested the significance of the overall model compared to a null model (only including the intercept and variables for which we wanted to control) to test whether or not main effects or their interaction overall had a significant effect. For sampling time, we combined the timepoints 6-12hrs to an early, 18-24hrs to a middle and 30-42hrs to a late ‘infection period’ in all analyses, except the time-resolved Cox Proportional Hazard Model. In all analyses of the worker pupae, we accounted for the effect of population (Jena, Seva), by including population also in the null model (since inclusion as random effect would require five levels)^32^. This was not required for the queen pupae, as they were only available from the Jena population. When multiple models were performed on related data, such as one model each for the four candidate CHCs for both worker and queen pupae separately (Tables S1a & b,S3a & b), or for the 14 non-candidate peaks of the queen pupae (Table S5), we corrected for multiple testing using the Benjamini-Hochberg procedure^33^ to protect against a false discovery rate (FDR) of 5%. When the overall model remained significant after correction for multiple testing, we proceeded to test the significance of interaction and, where applicable, the main effects, using likelihood ratio (LR) tests^34^. For models with significant interaction, the main effects cannot be reliably interpreted, and are hence not reported. All posthoc comparisons also included correction for multiple testing. Effect sizes were only calculated when a significant effect was found, using the packages ‘rstatix’ (vs 0.7.2)^35^ and ‘emmeans’ (vs 1.5.4)^36^. All reported p-values are exact unless <0.0001 and two-sided.

#### Pupal chemical compounds in dependence of treatment

To determine whether the pupae-produced amounts of the four candidate CHCs identified by Pull et al. 2018^9^ differed in dependence of infection status and presence/absence of workers – in particular, if the CHC abundances were different in infected pupae in the presence (I+W+) or absence (I+W-) of workers, and compared to the noninfected controls (I-W+, I-W-) – we ran a linear model per peak, testing for a significant interaction between our two main effects, thereby controlling for population (Jena and Seva) for the worker pupae and infection period (early, middle, late) for both worker and queen pupae by inclusion into the null model. Since we found no indication for signalling in the four candidate peaks, which Pull et al.^9^ had identified for worker pupae, in the queen pupae (Tables S3a,S4), we expanded our analysis for the queen pupae to the remaining 14 peaks (Table S5).

Prior to statistical analysis, we obtained normality in the worker pupae by transforming all four candidate peaks by the ordered quantile normalisation. For the queen pupae, we had to use different transformations for different CHCs, i.e. box-cox transformation for candidate compound C33:2 and non-candidates C33:1 (RI 3288), 13MeC33, log transformation for candidate compound C33:1 (RI 3279) and non-candidates 3MeC29, C30, C31, 3MeC33, square root transformation for candidate C35:2, ordered quantile normalisation for candidate C35:2+C35:1 and non-candidates C27, C28, C29, C33, C34, C35+13MeC35:1, C37 and asin transformation for non-candidate 3MeC33:1.

As we found that signalling was restricted to the two immune-associated compounds (C33:2 and C33:1) in the worker pupae, we wanted to test whether the combined differences of these two signalling compounds would lead to a significantly different chemical profile of the infected pupae in worker presence (I+W+) to the other groups. To this end, we carried out a combined analysis aggregating C33:2 and C33:1 into one analysis after having z-transformed each compound separately. We ran mixed linear models (LMM) with the same variables as above, yet in addition accounting for compound (C33:2, C33:1) in the null model. We further included sample ID as a random effect to control for non-independence of data points. After finding a significant interaction in the worker pupae (but no significance in the queen pupae, Table S4), we carried out all-pairwise posthoc tests for the worker pupae (Table S2) using the ‘emmeans’ package^36^.

To determine if individual pupal infection load affected the pupal amount of the two signalling compounds C33:2 and C33:1 in the worker pupae, we ran separate linear models (LM) for each compound, controlling for population, worker presence and infection period in the null model for the subset of the infected (I+) worker pupae (Table S1b). We did the same analysis for the infected (I+) queen pupae for consistency, except here not having to control for population (Table S3b).

#### Unpacking of pupae

Cox proportional-hazards regression analysis (‘survival’^37^ v.3.2.7) was used to determine, for both the worker and the queen pupae, whether unpacking occurred differently for the infected than the control pupae (worker pupae I+W+: n=133, I-W+: n=57; queen pupae: I+W+: n= 45, I-W+: n= 20), again accounting for ant population (Jena, Seva) in the worker pupae. We calculated the hazard ratio of how much more frequently unpacking occurred in the infected vs control pupae only for the worker pupae, as we found no significant difference in the queen pupae.

#### Infection progression

To assess how pathogen load progressed over the course of infection for the pupae of both castes, we ran separate linear models for worker and queen pupae, testing for the effect of infection period on the infection load (infected (I+) worker pupae: n=201, queen pupae: n=64). We controlled for worker presence in both worker and queen pupae models (Tables S7a &b) and as above, for population in the model for the worker pupae only (Table S7a). Following a significant effect of infection period on fungal pathogen load for both queen and worker pupae, we performed all-pairwise posthocs on whether the pupal infection loads differed between the early, middle and late stages of infection period using the ‘emmeans’ package^36^ (Tables S7a,b). Prior to analysis, we obtained normality of the data distribution by transforming data using the ordered quantile normalisation.

## Supporting information

Supplemental Tables S1-8, Figures S1-2, with suppl. references

## Acknowledgements

We thank Joergen Eilenberg and Nicolai V. Meyling for the fungal strain, the Social Immunity team at ISTA for ant collection and experimental help, Hanna Leitner and Jessica Kirchner for molecular support, Marco Ribezzi for advice on calculations and the ISTA Lab Support Facility, including the mass spectrometry unit, for general and chemical laboratory support. We thank Michaela Hönigsberger, Thomas Schmitt, Chris Pull and Michael Sixt for discussion and the Social Immunity team for comments on the manuscript.

## Author contributions

EHD and SC conceived the study; EHD, NK, JR, FS and AVG performed the experiments, NK generated the chemical and AVG the molecular data; EHD analysed and curated all data, performed the statistical analysis and made the figures after joint conceptualisation with SC; EHD and SC wrote the paper with input by NK and AVG; all authors approved the final manuscript version. Funding was obtained by SC.

