## Supplemental Tables S1-8, Figures S1-2, with suppl. references for "Altruistic disease signalling in ant colonies"

**Affiliations**

This file contains:

Supplementary Tables 1-8

Supplementary Figures 1-2

Supplementary References

### Supplementary Tables

**Supplementary Table 1. Effect of pupal infection and worker presence on worker pupae CHCs. (a)** Statistical results for each of the four CHCs identified by Pull et al 2018[1] as possible unpacking cue or signal in *L. neglectus* worker pupae, containing the two immune-associated compounds C33:2, and C33:1, as well as C35:2, and co-eluting C35:2 + C35:1. For the relative abundance of each peak, we used a Linear Model (LM) to test for a significant interaction between the main effects of pupal infection and the presence/absence of tending workers, if the overall model was significant. Sample size of total 323 worker pupae, of which n=69 infected without workers (I+W-), n=133 infected with workers (I+W+), n=64 uninfected controls without workers (I-W-) and n=57 uninfected controls with workers (I-W+)). **(b)** Following the significant interactions found for the two immune-associated compounds C33:2 and C33:1 we performed a linear model (LM) per compound to test for a significant effect of infection load on compound abundance, controlling for population, infection period and worker presence. The analysis focuses only on fungus-infected pupae (n=201, since PCR could not be conducted on one sample). For both (a) and (b), the data were transformed using the ordered quantile normalisation prior to analyses. Model details, compound ID (see also Table S8), test statistic, degrees of freedom (df), p-value and effect size given. Exact, two-sided p-values adjusted for multiple testing (as detailed in the Online Methods) are reported unless <0.0001. Significant p values highlighted in bold, "-" indicates non-applicable due to non-significance. Note for (b) that already the uncorrected raw p-values were >0.107, hence the absence of significance was not induced by correction for multiple testing.

**a)**

| **Relative compound abundance – worker pupae** | | | | | | | | |
|---|---|---|---|---|---|---|---|---|
| LM | compound | overall model | | | interaction | | | |
| | | $\chi^2$ | df | p-value | $\chi^2$ | df | p-value | Effect size ($\eta^2$p) |
| **Abundance ~ infection treatment * worker presence** + population + infection period | C33:2 | 32.812 | 3 | **<0.0001** | 7.044 | 1 | **0.0080** | 0.02 |
| | C33:1 | 17.248 | 3 | **0.0008** | 4.419 | 1 | **0.0356** | 0.01 |
| | C35:2 | 18.571 | 3 | **0.0007** | 0.763 | 1 | 0.382 | - |
| | C35:2+ C35:1 | 3.759 | 3 | 0.289 | - | - | - | - |

**b)**

| Effect of infection load on compound abundance – worker pupae | | | | |
|---|---|---|---|---|
| LM | compound | overall model | | |
| **Abundance ~ infection load** + population + infection period + worker presence | | $\chi^2$ | df | p-value |
| | C33:2 | 1.939 | 1 | 0.164 |
| | C33:1 | 2.598 | 1 | 0.164 |

**Supplementary Table 2. Statistical results of comparison of the four worker pupae groups based on their immune-associated CHCs.** Linear Mixed Model (LMM) results testing the differences between the four treatment groups, i.e. infected worker pupae without workers (I+W-), infected worker pupae with workers (I+W+), control worker pupae without infection and without workers (I-W-) and control worker pupae with workers (I-W+), based on both C33:2 and C33:1. Sample sizes as in Table S1. Data were transformed using the ordered quantile normalisation prior to analyses. Model details, test statistic, degrees of freedom (df) and p-value for the overall model and all-pairwise posthoc comparisons are given. Exact, two-sided p-values adjusted for multiple testing are reported. Significant p-values highlighted in bold and shown by letters a and b for the two significance groups (I+W+ vs all others) in Fig. 1A. "-" indicates non-applicable due to non-significance.

| Relative compound abundance of combined immune-associated compounds - worker pupae | | | | | | | | | | |
|---|---|---|---|---|---|---|---|---|---|---|
| LMM | compounds | overall model | | | interaction | | | pairwise comparisons | | |
| | | $\chi^2$ | df | p-value | $\chi^2$ | df | p-value | Pair | p-value | Effect size (Cohen's d) |
| **Abundance ~ infection treatment * worker presence** + compound + population + infection period + (1\|individual) | C33:2 and C33:1 | 16.554 | 3 | **0.00087** | 4.575 | 1 | **0.0324** | I+W+ vs I+W- | **0.0001** | 0.778 |
| | | | | | | | | I+W+ vs I-W+ | **0.012** | 0.527 |
| | | | | | | | | I+W+ vs I-W- | **0.002** | 0.623 |
| | | | | | | | | I+W- vs I-W- | 0.535 | - |
| | | | | | | | | I+W- vs I-W+ | 0.352 | - |
| | | | | | | | | I-W- vs I-W+ | 0.657 | - |

**Supplementary Table 3. Effect of pupal infection and worker presence on queen pupae CHCs. (a)** Statistical results for the four candidate CHCs for the queen pupae (as in Table S1a for the worker pupae). For none of the four peaks, its relative abundance was found to significantly differ in the overall LM, so that no further statistics were performed. Sample size of a total of 103 queen pupae, of which n=19 infected without workers (I+W-), n=45 infected with workers (I+W+), n=19 uninfected controls without workers (I-W-) and n=20 uninfected controls with workers (I-W+)). **(b)** As for the worker pupae (Table S1b) we also performed a linear model (LM) each to test for an effect of infection load on abundance of the two immune-associated compounds (C33:2 and C33:1), controlling for infection period and worker presence/absence. Analysis carried out on fungus-infected pupae only (n=64). To obtain normality of data distributions, data for each compound were transformed prior to analyses using the following transformations: C33:2 box-cox transformed, C33:1 log transformed, C35:2 square root transformed; C35:2+C35:1 ordered quantile normalisation, and infection load data were transformed using the ordered quantile normalisation. Model details, compound ID (see also Table S8), test statistic, degrees of freedom (df), and p-value given. Exact, two-sided p-values adjusted for multiple testing are reported. Note that also the uncorrected raw p-values were >0.472 in all of the compounds in (a) and >0.534 in (b), hence the absence of significance was not driven by multiple comparisons.

**a)**

| **Relative compound abundance – queen pupae** | | | | |
|---|---|---|---|---|
| LM | compound | overall model | | |
| | | $\chi^2$ | df | P-value |
| **Abundance ~ infection treatment * worker presence** + infection period | C33:2 | 1.789 | 3 | 0.816 |
| | C33:1 | 1.044 | 3 | 0.816 |
| | C35:2 | 2.516 | 3 | 0.816 |
| | C35:2+C35:1 | 0.940 | 3 | 0.816 |

**b)**

| **Effect of infection load on compound abundance – queen pupae** | | | | |
|---|---|---|---|---|
| LM | compound | overall model | | |
| | | $\chi^2$ | df | p-value |
| **Abundance ~ infection load** + infection period + worker presence | C33:2 | 0.0309 | 1 | 0.861 |
| | C33:1 | 0.386 | 1 | 0.861 |

**Supplementary Table 4. Statistical results of comparison of the four queen pupae groups based on their immune-associated CHCs.** Linear Mixed Model (LMM) results testing the differences between the four treatment groups, i.e. infected or uninfected queen pupae, each with or without workers, based on both C33:2 and C33:1. Sample sizes as in Table S3. To obtain normality, data were transformed using the Yeo-Johnson transformation. Model details, test statistic, degrees of freedom (df) and exact, two-sided p-values are given. Since the overall model was non-significant, indicated by ns in Fig. 1B, no pairwise posthoc comparisons were calculated.

| **Relative compound abundance of combined immune-associated compounds in the queen pupae** | | | | |
|---|---|---|---|---|
| LMM | compounds | overall model | | |
| | | $\chi^2$ | df | p-value |
| **Abundance ~ infection treatment * worker presence** + compound + infection period + (1\|individual) | C33:2 and C33:1 | 2.736 | 3 | 0.434 |

**Supplementary Table 5. Statistical testing for a potential signalling of queen pupae via the non-candidate CHCs**. To test whether queen pupae may use different compounds than the candidates identified by Pull et al. 2018[1] for the worker pupae for possible disease signalling, we also tested the remaining 14 compounds of the pupal chemical profile (for which quantification of their $^{12}C$ and $^{13}C$ proportions was possible for both worker and pupae; see Online Methods), for a possible difference according to infection treatment and worker presence. Sample sizes as in Table S3. To obtain normality of data distributions, data for each compound were transformed prior to analyses using the following transformations: C27, C28, C29, C33, C34, C35+13MeC35:1, C37 ordered quantile normalisation; 3MeC29, C30, C31, 3MeC33 log transformation; C33:1 (non-candidate C33:1 with RI 3288; Table S8), 13MeC33 box-cox transformation; 3MeC33:1 asin transformation. Model details, compound ID (see also Table S8), test statistic, degrees of freedom (df) and exact, two-sided p-value after correction for multiple testing (as detailed in the Online Methods) given. None of the overall models were significant after p-value adjustment for multiple testing. Note that three compounds (C27, 13MeC33 and 3MeC33) were significant in the overall model in the absence of adjustment, yet their interaction terms were all >0.43, therefore supporting that we found no evidence of chemical signalling in the queen pupae.

| Relative compound abundance of non-candidate compounds in queen pupal profile | | | | |
|---|---|---|---|---|
| LM | compound | overall model | | |
| **Abundance ~ infection treatment * worker presence** + infection period | | $\chi^2$ | df | p-value |
| | C27 | 9.74 | 3 | 0.141 |
| | C28 | 4.38 | 3 | 0.474 |
| | C29 | 3.202 | 3 | 0.474 |
| | 3MeC29 | 5.100 | 3 | 0.461 |
| | C30 | 3.596 | 3 | 0.474 |
| | C31 | 2.703 | 3 | 0.474 |
| | C33:1 | 1.988 | 3 | 0.575 |
| | C33 | 3.560 | 3 | 0.474 |
| | 13MeC33 | 8.935 | 3 | 0.141 |
| | 3MeC33:1 | 5.759 | 3 | 0.434 |
| | 3MeC33 | 10.908 | 3 | 0.141 |
| | C34 | 3.764 | 3 | 0.474 |
| | C35+13MeC35:1 | 3.067 | 3 | 0.474 |
| | C37 | 2.792 | 3 | 0.474 |

**Supplementary Table 6. Worker unpacking behaviour directed to infected vs uninfected pupae.** Statistical results for the cox proportional-hazards regression of the effect of infection on unpacking behaviour. Infected worker pupae elicited significantly more unpacking by the workers than uninfected controls, whilst there was no significant effect of infection treatment in the queen pupae. Model, test statistic, degrees of freedom (df), and p-value given. For the queen pupae, the overall model directly gives the effect of treatment, as all stem from the same population. For the significant effect of treatment (infected, uninfected) in the worker pupae, the hazard ratio of being unpacked when infected and its CI are reported ("-" for queen pupae due to non-significance). Significant p-values in bold. Samples sizes as in Table S1a,S3a.

| **Worker unpacking towards worker resp. queen pupae** | | | | | | | |
|---|---|---|---|---|---|---|---|
| Pupae | Cox proportional-hazards regression | overall model | | | infection treatment | | |
| | | Wald Test | df | p-value | p-value | HR | CI |
| **Worker** | **Unpacking ~ infection treatment** + population | 8.82 | 2 | **0.010** | **0.0037** | 4.542 | 1.64-12.60 |
| **Queen** | **Unpacking ~ infection treatment** | 2.49 | 1 | 0.115 | equal to overall | - | - |

**Supplementary Table 7. Infection load timeline in (a) worker pupae and (b) queen pupae.** Statistical results for the comparison in fungal infection loads of (a) worker pupae and (b) queen pupae at the different periods in infection progression from the start of the experiment (which began three days after pathogen exposure). Samples for the 'early' infection period were collected at 6 & 12 hours, for the 'middle' at 18 & 24 hours and for the 'late' at 30, 36 & 42 hours after the start of the experiment. Sample sizes for the worker pupae were n=75 early, n=57 middle and n=69 late infection period samples, whilst n=16 early, n=16 middle and n=32 late for the queen pupae. To obtain normality of data distribution, data were transformed prior to analyses using the ordered quantile normalisation. Model details, test statistic, degrees of freedom and p-value given. Pairwise tests between infection periods were corrected for multiple testing; effect sizes given for significant effects. "-" indicates non-applicable due to non-significance. Exact p-values reported unless <0.0001. Significant p-values in bold.

**a)**

| Infection progress – worker pupae | | | | | | |
|---|---|---|---|---|---|---|
| LM | overall model | | | pairwise comparisons | | |
| | $\chi^2$ | df | p-value | pair | p-value | effect size (Cohen's d) |
| **Infection load ~ infection period** + population + worker presence | 36.997 | 2 | **<0.0001** | early to middle | **0.0157** | 0.441 |
| | | | | middle to late | **0.0007** | 0.643 |
| | | | | early to late | **<0.0001** | 1.084 |

**b)**

| Infection progress – queen pupae | | | | | | |
|---|---|---|---|---|---|---|
| LM | overall model | | | pairwise comparisons | | |
| | $\chi^2$ | df | p-value | pair | p-value | effect size (Cohen's d) |
| **Infection load ~ infection period** + worker presence | 10.178 | 2 | **0.006** | early to middle | 0.161 | - |
| | | | | middle to late | **0.006** | -1.002 |
| | | | | early to late | 0.180 | - |

**Supplementary Table 8. Cuticular hydrocarbons (CHCs) in *Lasius neglectus* worker and queen pupae.**

Compound name (ordered by retention time), chain length and structure, as well as Kováts retention index (RI) of the 18 pupal cuticular hydrocarbons (CHCs), for which quantification of their $^{12}C$ and $^{13}C$ proportions was possible for both worker and pupae (see Online Methods). Compound identification as in Ugelvig et al. 2008 and Pull et al. 2018[1,2]. Peaks in bold have been identified as being upregulated in worker pupae infected with the fungal pathogen *Metarhizium brunneum* by Pull et al. 2018. All compounds are present in both worker and queen pupae.

| **Compound name** | **Chain length and structure** | **RI** |
|---|---|---|
| *n*-Heptacosane | C27 | 2699 |
| *n*-Octacosane | C28 | 2799 |
| *n*-Nonacosane | C29 | 2902 |
| 3-Methylnonacosane | 3MeC29 | 2974 |
| *n*-Triacontane | C30 | 2999 |
| *n*-Hentriacontane | C31 | 3100 |
| **Tritriacontadiene** | **C33:2** | **3251** |
| **Tritriacontene** | **C33:1** | **3279** |
| Tritriacontene | C33:1 | 3288 |
| *n*-Tritriacontane | C33 | 3300 |
| 13-Methyltritriacontane | 13MeC33 | 3326 |
| 3-Methyltritriacontene | 3MeC33:1 | 3353 |
| 3-Methyltritriacontane | 3MeC33 | 3373 |
| *n*-Tetratriacontane | C34 | 3402 |
| **Pentatriacontadiene** | **C35:2** | **3447** |
| **Pentatriacontadiene + Pentatriacontene** | **C35:2 + C35:1** | **3475** |
| *n*-Pentatriacontane + 13-Methylpentatriacontene | C35+13MeC35:1 | 3500 |
| *n*-Heptatriacontane | C37 | 3697 |

**Supplementary Figures**

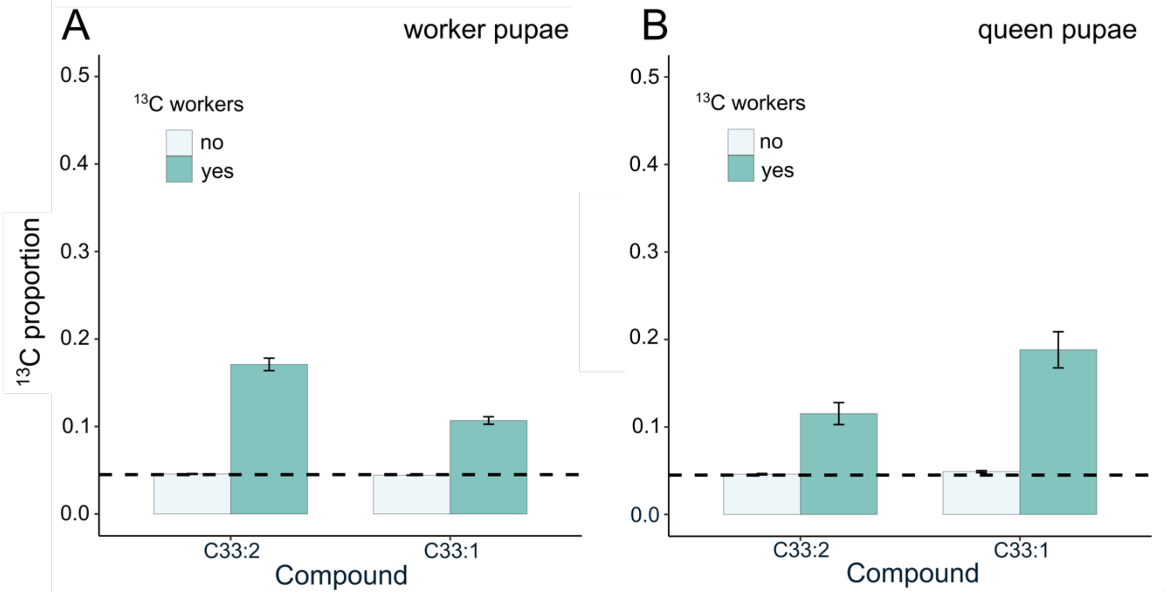


**Supplementary figure 1) $^{13}$C transfer to worker and queen pupae.** The proportion of $^{13}$C relative to the total amount of carbon ($^{12}$C and $^{13}$C combined) measured in the two immune-associated compounds C33:2 and C33:1 of the extracts from of (A) worker and (B) queen pupae kept in the presence of $^{13}$C-enriched workers (darker green bars; worker pupae n=190; queen pupae n=65) vs alone (light green bars; worker pupae n=133, of which in 5 pupae the C33:2 proportions could not be calculated due to too low compound abundance; queen pupae n=38). The accompanying workers integrated $^{13}$C into their CHCs during the three-week feeding period, leading to values clearly increased above the average natural 0.044 value (C33:2 mean: 0.475 ± 0.009 sem, C33:1 mean: 0.455 ± 0.013 sem; n=255 pools of 2 workers each, total 510 workers). The pupae kept with these enriched workers also show elevated levels of $^{13}$C compared to the natural level of 0.044 (dashed line), whilst the pupae kept alone do not. Bars show means, error bars depict ± sem.

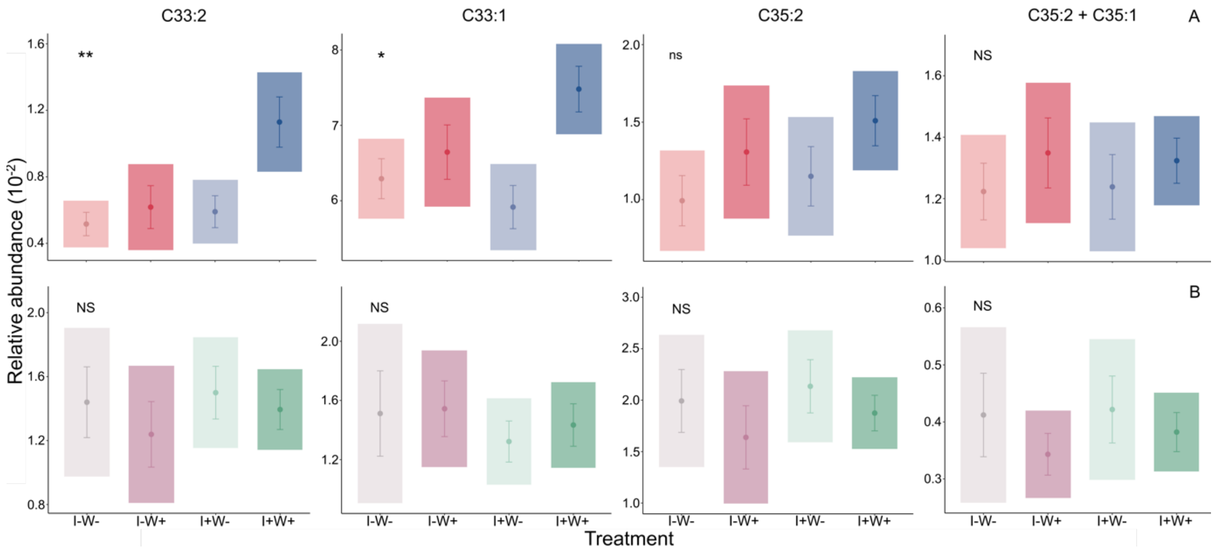


**Supplementary figure 2) CHC abundance separately for the four possible candidate peaks** identified by Pull et al. 2018. Individual graphs showing the relative abundance of the four candidate CHC peaks consisting of compounds tritriacontadiene (C33:2), tritriacontene, (C33:1), pentatriacontadiene, (C35:2), and co-eluting pentatriacontadiene and pentatriacontene, (C35:2 + C35:1) for (A) worker and (B) queen pupae not infected and kept without workers (I-W-), not infected and kept with workers (I-W+), infected and kept without workers (I+W-) and infected kept with workers (I+W+). Worker pupae shown in blue for infection and red for control treatment, and queen pupae in green, resp. purple; absence of workers indicated by pale colours. Dots and bars represent mean ± sem, while shaded area represents 95% confidence intervals of the mean. Statistics for worker pupae provided in Table S1, for queen pupae in Table S3; significant interaction between infection treatment and worker presence shown by * for p<0.05 and ** for p<0.001 and ns for p>0.05 (worker pupae C35:2); NS represents non-significant overall model (worker pupae C35:2+C35:1 and all queen pupae peaks).
